# Motorized shoes induce robust sensorimotor adaptation in walking

**DOI:** 10.1101/788943

**Authors:** Yashar Aucie, Xunjie Zhang, Randy Sargent, Gelsy Torres-Oviedo

## Abstract

The motor system has the flexibility to update motor plans according to systematic changes in the environment or the body. This capacity is studied in the laboratory through sensorimotor adaptation paradigms imposing sustained and predictable motor demands specific to the task at hand. However, these studies are tied to the laboratory setting. Thus, we asked if a portable device could be used to elicit locomotor adaptation outside the laboratory. To this end we tested the extent to which a pair of motorized shoes could induce similar locomotor adaptation to split-belt walking, which is a well-established sensorimotor adaptation paradigm in locomotion. We specifically compared two groups of young, healthy subjects adapted on the treadmill by moving their feet at different speeds with a split-belt treadmill or with a pair of motorized shoes. We found that the adaptation of joint motions and measures of spatial and temporal asymmetry, which are commonly used to quantify sensorimotor adaptation in locomotion, were indistinguishable between groups. We only found small differences in the join angle kinematics during baseline walking between the groups-potentially due to the relatively large weight and height of the motorized shoes. Our results indicate that robust sensorimotor adaptation in walking can be induced with a paired of motorized shoes, opening the exciting possibility to study sensorimotor adaptation during more realistic situations outside the laboratory.

## 1 Introduction

The motor system has the flexibility to update motor plans according to systematic changes in the environment or the body. This human ability is studied in the laboratory through sensorimotor adaptation paradigms imposing sustained and predictable motor demands specific to the task at hand, such as unusual visuomotor rotations (e.g., (Krakauer et al., 2000) or constant forces during walking (Savin et al., 2010) or reaching (Shadmehr and Mussa-ivaldi, 1994). For example, split-belt walking is a well-established paradigm in which subjects update spatiotemporal gait features in response to a persistent speed difference between their legs (Malone et al., 2012). Important motor adaptation principles have been learned from these sensorimotor adaptation paradigms, such as the computations underlying motor adaptation (Haruno et al., 2001; Smith et al., 2006; Thoroughman and Shadmehr, 2000) or neural structures involved in this process (Deuschl et al., 1996; Morton and Bastian, 2006; Smith and Shadmehr, 2005). However, there are inherent limitations to laboratory-based studies that bring into question the extent to which principles governing motor adaptation apply to motor learning in the real-world.

Specifically, there are task-constraints in laboratory-based studies that limit our ability to investigate factors that are critical for motor learning outside the laboratory setting. For example, laboratory-based protocols are designed such that new motor behaviors are acquired quickly. While this enables to characterize the evolution of the adaptation process from transient to steady-state behaviors (Smith et al., 2006), it limits our understanding of practice, which is critical for mastering any real-world skill (Ericsson and Pool, 2016; Haith and Krakauer, 2018). Further, we constrain movements by for example making people walk at a constant speed (Dietz et al., 1994), or reach to a certain direction (Krakauer et al., 2000). This is done to simplify the control variables affecting the studied behavior, and at the extreme this could yield to the study of unnatural behaviors, which principles might not apply to realistic situations. A byproduct from task-constraints is the context-specificity of motor patterns learned in the laboratory –that is movements adapted with the device do not carry over when moving without the device (Kluzik et al., 2008; Torres-Oviedo and Bastian, 2010). This is detrimental not only because it limits our capacity for studying the generalization of motor learning across distinct situations, but also because it limits the possibility for using laboratory-based tasks for motor rehabilitation. Notably, it is well-accepted that the generalization of motor patterns from trained to untrained situations can be improved when the two contexts are more similar to one another (Bouton et al., 1999; Spear, 1978; Tulving and Thomson, 1973). Thus, there could be more generalization of laboratory-based knowledge to realistic situations when the tasks studied in the laboratory are more similar to those observed under naturalistic conditions.

Portable devices may offer the possibility to overcome the limitations of laboratory-based studies of motor learning. For example, portable devices allow us to investigate motor learning in real-life settings, such as studies of surgical training with the same tools that are used at the clinic (Sharon et al., 2017). In addition, the portability of training devices also enables the study of extended practice since individuals are not constrained to only train in the laboratory setting (Hardwick et al., 2019). Further, portable devices might allow for more complex movements that involve the whole body (Haar et al., 2019), which might lead to greater motor variability –a key factor for motor learning (Kelly and Sober, 2014; Therrien et al., 2016; Wu et al., 2014). In the context of locomotion there have been efforts to develop portable devices to study motor adaptation (Handzic et al., 2011; Handzic and Reed, 2013; Lahiff et al., 2016). However, gait adjustments induced by these devices are not as robust as the ones observed with laboratory-based apparatus such as split-belt treadmills. Thus, we asked if a pair of motorized shoes could induce locomotor adaptation comparable to split-belt walking, which is a well-established sensorimotor adaptation paradigm in locomotion.

We specifically hypothesized that introducing a speed difference between subject’s feet with the motorized shoes would result in adaptation of spatiotemporal gait patterns similar to split-belt walking. To test this hypothesis, we compared locomotor adaptation on motorized shoes vs. the split-belt to comparable speed differences imposed by the two devices. If the locomotor adaptation with the motorized shoes is as robust as the one observed with the split-belt walking paradigm, participants could start wearing these shoes outside the laboratory, which would offer the exciting possibility to study locomotor learning under more realistic situations.

## 2 Methods

### 2.1 Participants

We investigated if a pair of motorized shoes could induce locomotor adaptation and after-effects similar to a split-belt treadmill. To this end, a group of 18 young and healthy adults were adapted using either 1) the motorized shoes that imposed speed differences between the feet using actuated wheels under the shoe (Motorized shoes group: n=9; 3 females: 26.6 ± 3.5 years) or 2) a split-belt treadmill, in which belts moved at different speeds (Split-belt group: n=9; 4 females: 25.3 ± 4.3 years). The Institutional Review Board at the University of Pittsburgh approved our experimental protocol and all participants gave their written informed consent before being tested.

### 2.2 Set up

The Motorized shoes group walked on the treadmill while wearing the custom made motorized shoes (Nimbus Robotics, Pittsburgh PA) as shown in Figure 1A on top of their normal walking shoes. In brief, the shoes were designed to move an individual (weighing less than 100 Kg) up to 1 m/s forward. Each of the motorized shoe consisted of a motor, a controller box, a gearbox, two toothed timing belts, and 4 rubber wheels (Figure 1B). Lithium batteries (3V) were used to power the motor, which rotated the timing belts via a gearbox connecting the two. The timing belts and rubber wheels were coupled to rotate the wheels such that they locked the non-actuated shoe during stance (∼0 m/s) and moved the actuated shoe forward at a linear speed of 1 m/s. The controller boxes received signals through a remote controller operated by the experimenter. All the software for the controller boxes and the remote controller were written with Python. Details on the control software are published in (Zhang, 2017) and a detailed description of the motorized shoes will be revealed in the full utility patent (currently in provisional status). The Split-belt group did not wear the motorized shoes and walked with their regular shoes on an instrumented split-belt treadmill (Bertec, Columbus Ohio).

**Figure 1.**
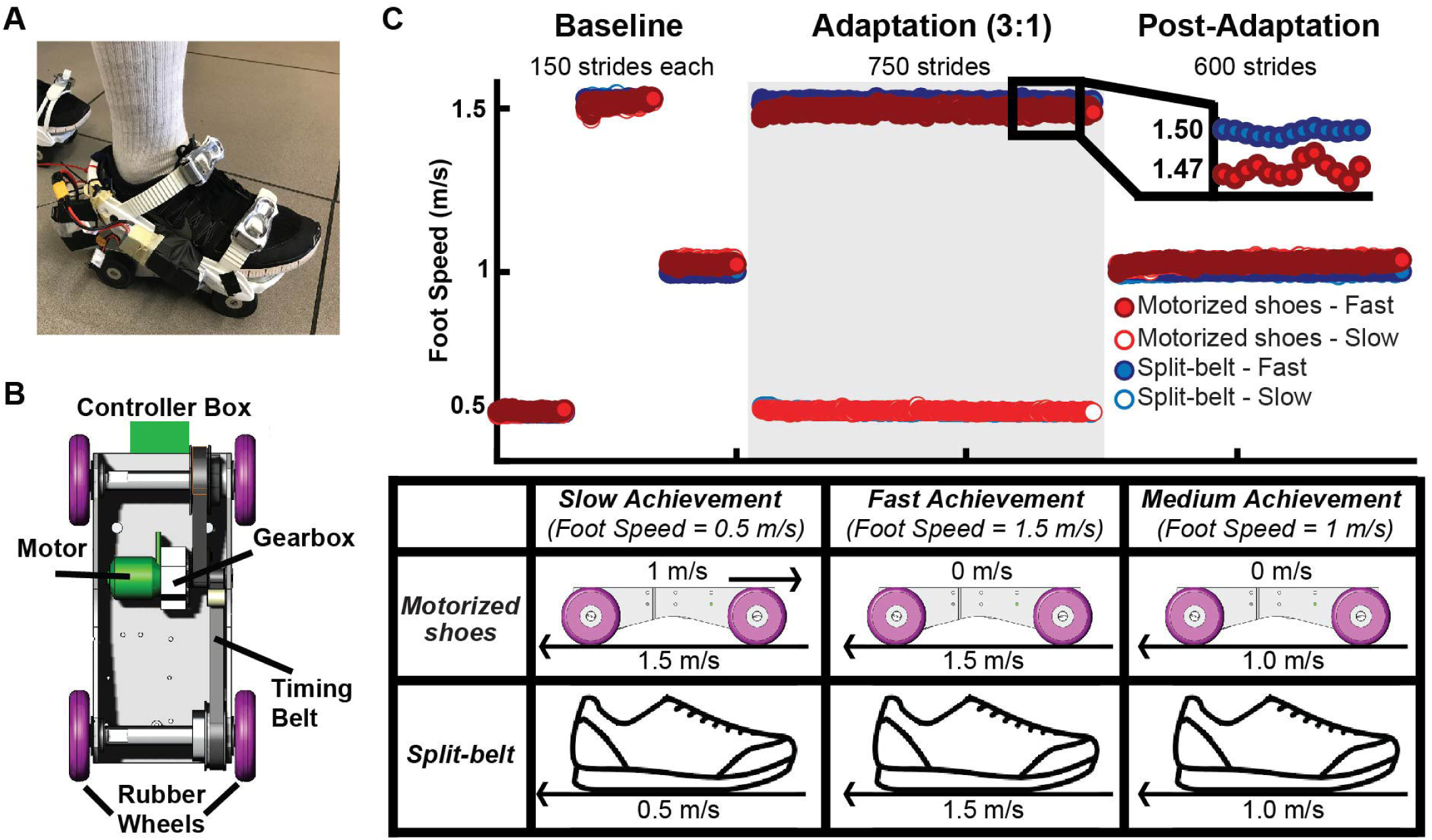
A) A motorized shoe involving proprietary technology was used to induce adaptation in the Motorized shoes group. B) Motorized shoes’ design schematic that is consists of a motor, a controller box, a gearbox, two toothed timing belts, and four rubber wheels. C) Mean time courses for foot speed across subjects for the Motorized shoes and the Split-belt groups. The white background indicates experimental epochs of ‘tied’ walking when both feet moved at the same speed, whereas the grey background indicates the epoch of ‘split’ walking when the dominant leg moved 3 times faster than the non-dominant leg. The table summarizes the procedure used to set the slow, fast, and medium speeds for each foot. Same procedure was used in all epochs. It is worth pointing out that the treadmill always moved at 1.5 m/s during adaptation in the Motorized shoes group. The speed difference between feet was achieve by locking the wheels on the fast side and moving the slow foot forward at 1 m/s to obtain a net speed of 0.5 m/s on the slow side. Of note, the foot’s speed on the fast side was slightly slower on the Motorized shoes than the Split-belt group.

### 2.3 General Paradigm

All subjects adapted following a conventional sensorimotor adaptation paradigm that consisted of three walking conditions: baseline, adaptation, and post-adaptation (Figure 1C - Top). During these periods subjects’ feet moved at one of three possible speeds: slow (0.5 m/s), medium (1m/s), or fast (1.5 m/s). The speeds at which each foot moved in the Split-belt group was set by the belt speed under each foot, whereas in the motorized shoes group were set by the combined effect of the actuated shoes and the belts’ speed (Table displayed on Figure 1C - Bottom). In particular, the slow foot speed was achieved by activating the motorized shoe that moved the foot in contact with the ground forward at 1 m/s, while the treadmill moved it backwards at 1.5 m/s such that the net foot speed was 0.5 m/s. On the other hand, the medium and fast foot speed was achieved by locking the motorized shoes’ wheels such that the foot in contact with the ground moved at the speed of the belt (i.e., 1m/s or 1.5 m/s). This approach enabled us to move subjects’ feet in the Motorized shoes group at the same speeds as in the Split-belt group, while both belts moved at the same speed as in a regular treadmill. We had so many different possibilities to achieve the net speed and we chose the one to maximize the duration of experiment for a given battery life. A baseline period was collected during which both feet moved at either slow, fast, or medium speeds for 150 strides each (Figure 1C - Top). The baseline behavior during the slow and fast speeds served as a reference for the adaptation condition when the feet moved at different speeds, whereas the medium speed served as a reference for the post-adaptation period when the two feet move at the same medium speed. Importantly, the baseline behavior was matched not only in the speed at which the feet moved, but also on how this speed was achieved. In other words, the Motorized shoes group wore the motorized shoes during the entire duration of the experiment and the speed was regulated as indicated in the table on Figure 1C during baseline, adaptation, and post-adaptation. In the adaptation period, the dominant leg (self-reported leg to kick a ball) moved at 1.5 m/s (i.e., fast side) and the non-dominant leg moved at 0.5 m/s (i.e., slow side) for 750 strides (approx. 15 min). The speed difference and period duration was selected to match other split-belt walking studies showing robust gait adaptation (Sombric et al., 2019). Following the adaptation block, all participants experienced a post-adaptation period of 600 strides during which both feet moved at 1 m/s, which was the average speed of the fast and slow feet. The purpose of this phase was to measure the adaptation effects and its washout when the speed perturbation induced by different devices was removed.

### 2.4 Data Collection

All subjects walked on an instrumented treadmill either with or without the motorized shoes, while kinematic and kinetic data were collected to characterize subjects’ gait. Kinematic data were collected at 100 Hz with a passive motion capture system (Vicon Motion Systems, Oxford UK) and kinetic data were collected at 1000 Hz using force plates embedded in the treadmill. Gaps in raw kinematic data due to marker occlusion were filled by visual inspection of each subject in Vicon Nexus software. Positions from the toe, ankle (lateral malleolus), knee (lateral epicondyles) and the hip (greater trochanter) were collected bilaterally (Figure 2B). Heel-strikes (i.e., foot landing) and toe-offs (i.e., foot lift off) were identified using the normal ground reaction force (Fz). More specifically, heel-strike was defined as the instance when Fz> 30 N and toe-off as the instance when Fz < 30 N. We used this force threshold to have equivalent event detection (i.e., heel strike, toe off) on the treadmill for both groups since each of the motorized shoe weighted 17 N (∼1.7 kg in mass).

**Figure 2.**
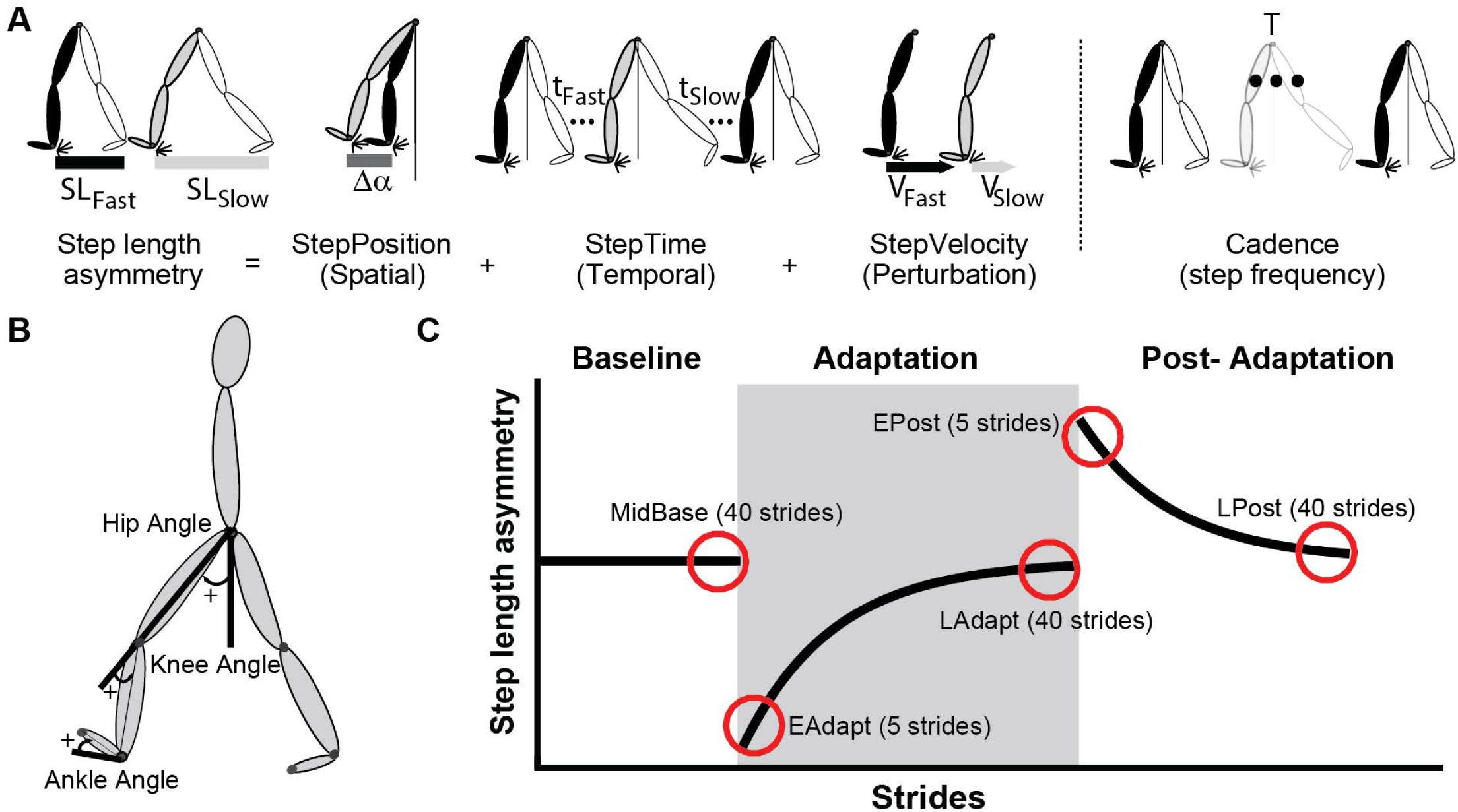
A) This schematic, adapted from (Finley et al., 2015), illustrates Step length asymmetry (defined as the difference between fast and slow step lengths, normalized by stride length), StepPosition, StepTime, StepVelocity, and Cadence parameters. B) Illustration of reflective marker positions and joint angle conventions. C) Epochs of interest are illustrated by the red circles placed over a schematic of step length asymmetry. Shaded gray area represents the adaptation period when the feet move at different speeds (“split” walking). “Tied” walking, when two legs are moving at the same speed, is indicated by the white regions.

### 2.5. Data Analysis

We compared the gait pattern between the Motorized shoes and Split-belt groups in terms of spatial and temporal symmetry measures that are known to adapt on the split-belt treadmill (Figure 2A) (Finley et al., 2015). Specifically, we used step length asymmetry as a robust measure of adaptation. Step length asymmetry was defined as the difference between step lengths (i.e., distance between ankles) when taking a step with the leg walking slow vs. the leg walking fast (Eq. 1). A zero value of step length asymmetry indicated that both step lengths were equal and a positive value indicated that the step length of the fast (dominant) leg was longer than the slow (non-dominant) leg. Step length asymmetry was further decomposed into StepPosition, StepTime, and StepVelocity because these parameters have been shown to be adapted differently during split-belt walking (Finley et al., 2015). The StepPosition quantified the difference in positions of the leading leg (i.e., leg in front of the body) between two consecutive steps (Eq. 2). The StepTime quantified the difference in the duration of each of these steps (Eq. 3). Lastly, the StepVelocity quantified the difference in the velocities of each foot with respect to the body for these two steps (Eq. 4). Since subjects take steps with different sizes, we normalized the differences in step length, StepPosition, StepTime and StepVelocity by their stride length, quantified as the sum of two step lengths. This allowed us to avoid intersubjective variability. For visualization purposes, these parameters were smoothed with a 5-step running average.

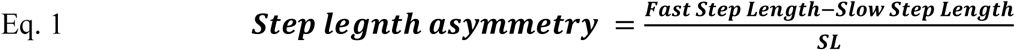

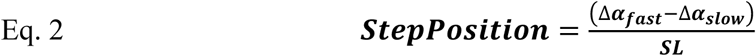

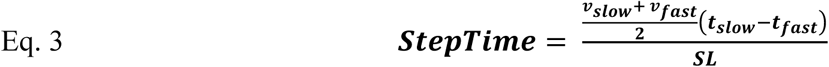

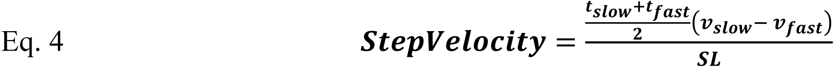

In these equations, Δ*α* indicates the difference between each foot’s position (i.e. ankle marker) and the body (i.e., mean position of the two hip markers) at ipsilateral heel strike (Figure 2A); In addition, *t* indicates the step time defined as the duration between the heel-strike of ipsilateral leg to the contralateral leg; and v indicates the step velocity quantified as the relative velocity of the foot with respect to the body. When walking on the treadmill, v*slow* and v*fast* approximated the speeds of the slow and fast belt, respectively. Therefore, StepVelocity was mostly reflective of belt speed difference, rather than subjects’ behavior. Finally, note that all measures were normalized by each subject’s stride length (SL, sum of both step lengths) to account for inter-subject differences in step sizes.

We also computed joint angles and cadence to determine the impact of the shoes on each foot’s motion and step frequency. Ankle, knee, and hip angles were computed on the sagittal plane (2D) since walking has a unique pattern of movement on that plane (Reisman et al., 2005). Joint angles were calculated such that flexion/dorsiflexion was positive and extension/plantarflexion was negative (Figure 2B). We also defined all angles to have value of 0° at the neutral standing position (i.e., full extension for knee and hip and a 90° degree angle between shank and foot for the ankle). More specifically, ankle angles were calculated as the angle between the foot (ankle marker to toe marker vector) and the shank (ankle marker to knee marker vector) subtracted from 90°. Knee angles were calculated as the angle between the shank and the thigh (knee marker to hip marker vector) subtracted from 180°. Lastly, we computed the hip angles as the angle between the thigh and the vertical unit vector. Angle data was time-aligned and binned to compute mean angle values over 6 intervals of interest during the gait cycle. This was done to focus on changes in angles within the gait cycle, rather than on changes due to differences in cycle duration across the distinct walking conditions (Dietz et al., 1994; Reisman et al., 2005). More specifically, we computed averaged angle values over 6 phases of interest: Double support (DS1, DS2), Single stance (SS1, SS2), and the swing phases (SW1, SW2). Double support during early stance (DS1) was defined as the period from heel strike to contralateral toe off. Single stance (from contralateral toe-off to contralateral heel strike) was divided into 2 equal phases (SS1, SS2). Double support during late stance (DS2) was defined as the interval from contralateral heel strike to ipsilateral toe off. Finally, the swing phase (from ipsilateral toe-off to ipsilateral heel-strike) was divided into 2 equal phases (SW1, SW2). Joint angles were assessed in 8 subjects per group since the remaining 2 subjects (one per group) was missing essential marker data. Lastly, we computed cadence (i.e. number of strides per second) to determine if this gait feature was altered by wearing the motorized shoes.

### 2.6 Outcome measures

Each gait parameter was analyzed during four experimental epochs of interest to compare the adaptation and after-effects between the Motorized shoes and the Split-belt treadmill groups. The epochs of interest included: early adaptation (EAdapt, first 5 strides), late adaptation (LAdapt, last 40 strides), early post-adaptation (EPost, first 5 strides), and late post-adaptation (LPost, last 40 strides) (Figure 2C). All of the parameters were corrected by any baseline biases (MidBase, last 40 strides. EAdapt gave us information about the induced perturbation by the ‘split’ condition, while the LAdapt provided information regarding the steady-state behavior at the end of the adaptation trial. The behavior during EPost was quantified to assess how much the subjects adapted to the new walking pattern (e.g., after-effects). Finally, we assessed LPost behavior to ensure that the subjects returned to their baseline walking behavior (e.g., washout). Moreover, we used joint angle measures to determine the effect of the motorized shoes on the overall gait pattern. The 6-joint parameters for each angle were computed for the following epochs of interest: slow baseline (SBase), fast baseline (FBase), medium baseline (MidBase) and the late adaptation (LAdapt). To this end we computed the averaged value over the last 40 strides for each one of the 4 experimental epochs of interest (i.e., SBase, FBase, MidBase, and LAdapt). The five strides at the beginning and end of each trial were discarded to eliminate effects of starting and stopping of the treadmill.

### 2.7 Statistical Analysis

Separate two-way repeated measures ANOVAs were used to test the effects of epochs (i.e., EAdapt, LAdapt, EPost, and LPost) and groups (i.e., Motorized shoes vs. Split-belt) on each of our gait parameters (i.e., Step length asymmetry, Step lengths, StepPosition, StepTime, StepVelocity, and Cadence). Statistical analysis were done with unbiased data (i.e., MidBase was subtracted from all the epochs) to focus on changes that occurred beyond those due to distinct group biases. In case of significant main or interaction effects, we used Fisher’s post-hoc testing to determine whether values were different between groups. We chose this post-hoc testing to increase the false positive (Type I error); therefore, becoming more sensitive to potential group differences. Lastly, we performed a one-sided one sample t-test to determine whether early post-adaptation values were different from zero.

Two sets of correlations were performed to assess the association between StepVelocity and 1) StepPosition and StepTime in late adaptation and 2) StepPosition and StepTime after-effects. This was done because we observed speed differences between the groups (Figure 1C - Top) that could impact the extent of adaptation and after-effects on spatial and temporal measures.

Joint angles were compared across groups using unpaired t-test for each of the gait phases. We reasoned this was an appropriate statistical test to compare the behavior across groups given that joint angles are highly temporally correlated within the gait cycle and spatially correlated across segments. We subsequently corrected the significance threshold for each epoch using a Benjamini-Hochberg procedure (Benjamini and Hochberg, 1995), setting a false discovery rate of 5% (FDR correction). The reason for choosing this correction was due to higher number of comparisons that we made.

A significance level of α=0.05 was used for all statistical tests. Stata (StataCorp., Collage Station, TX, was used to perform the ANOVAs, whereas MATLAB (The MathWorks, Inc., Natick, MA, United States) was used for all other analyses.

## 3 Results

### Motorized shoes can induce robust sensorimotor adaptation of locomotion

Our results show that the motorized shoes were able to induce similar adaptation of step length asymmetry compared to the split-belt treadmill. Specifically, there were no significant group (F_(1,48)_=0.21, p=0.65) or group by epoch interaction effects (F_(3,48)_=1.26, p=0.29) on the adaptation of step length asymmetry, indicating that this parameter was similarly modulated throughout the experiment between the Motorized shoes and Split-belt groups (Figure 3A). We also observed a significant main effect of epoch (F_(3,48)_=94.91, p<0.001) and found that both groups had significant after-effects (Motorized shoes: p <0.001; Split-belt: p<0.001; Figure 3A). While modulation of step length asymmetry was indistinguishable between groups, we observed subtle differences in the adaptation of the fast leg’s step length. Specifically, we found a group by epoch interaction effect (F_(3,48)_=3.18, p=0.032; Figure 3B) driven by between-group differences during the early adaptation phase (p=0.012). Moreover, after-effects in this parameter were significant in the Motorized shoes group (p = 0.013), but not in the Split-belt group (p = 0.15). In contrast, the adaptation of the slow leg’s step length was similar across groups throughout the experiment (group: F_(1,48)_=0.63, p=0.44; group by epoch interaction: F_(3,48)_=0.69, p=0.49; Figure 3C). We only found a significant epoch effect on slow step length (F_(3,48)_=70.47, p<0.001) and substantial after-effects in both groups (Motorized shoes: p<0.001; Split-belt: p<0.001). In summary, the fast leg’s step length exhibited some adaptation differences between the Motorized shoes and Split-belt groups, but the overall adaptation of step length asymmetry was similar across groups.

**Figure 3.**
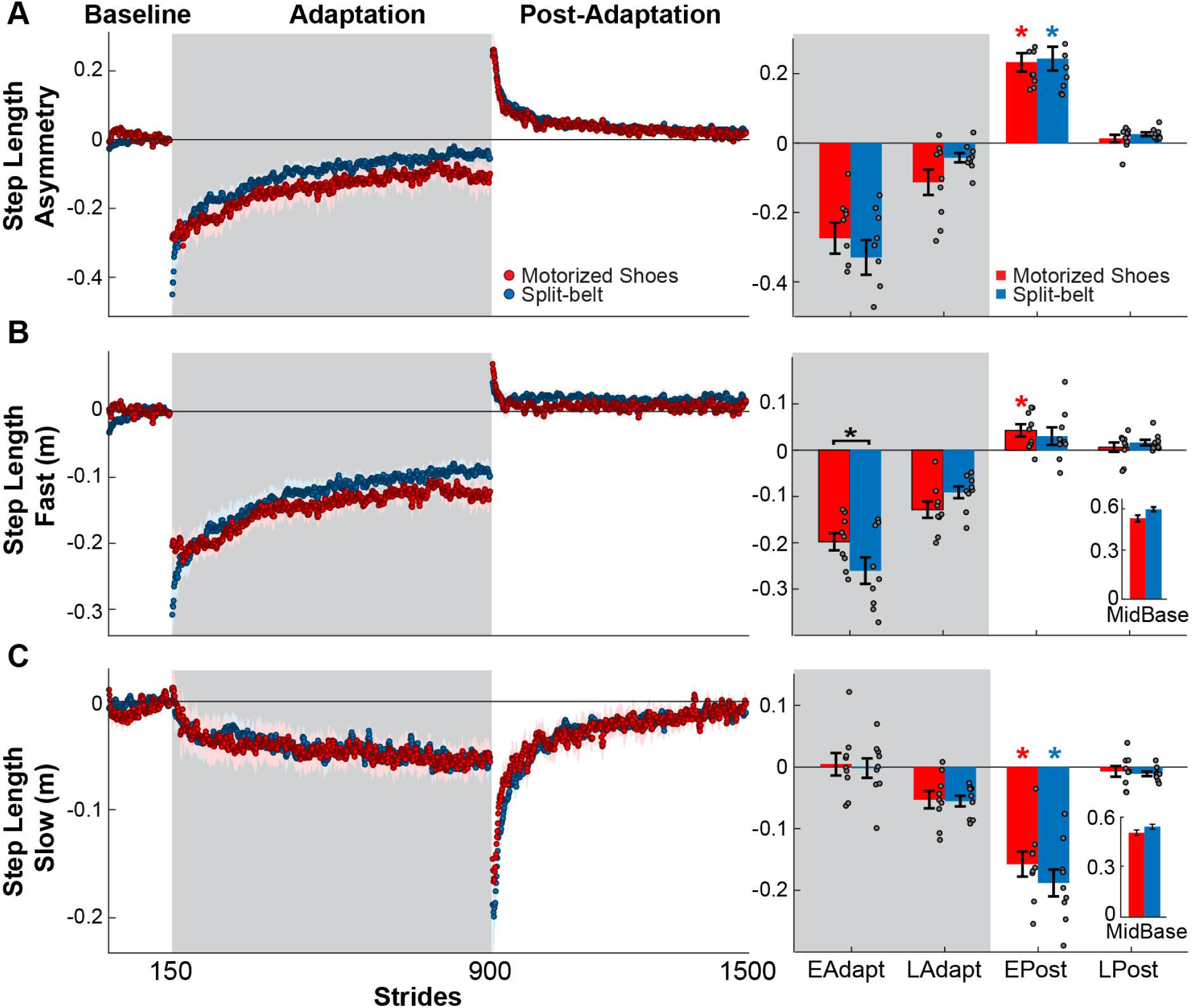
Modulation of step length asymmetry and step lengths. (A, B, C- Left Panel) Time courses for step length asymmetry and individual step lengths during medium baseline, adaptation and post-adaptation. Shaded gray areas represent the adaptation period during which the legs are moved at different speeds. “Tied” walking, both feet moved at the same speed, is indicated by the white regions. Colored dots represent the group average of 5 consecutive strides and colored shaded regions indicate the standard error for each group (Motorized shoes: red; Split-belt: blue). (A, B, C- Right Panel) Bar plots indicate the mean ± standard errors for step length asymmetry and step lengths for each group and epoch of interest. Note that the values were corrected for baseline biases. Significant differences for post-hoc tests were indicated as follows. Black asterisks over the bracket above each epoch represent statistical significant differences between the Motorized shoes and the Split-belt groups (p<0.05). Colored asterisks over the bars indicate significant after-effects (i.e., early post-adaptation is significantly different from baseline; p<0.05) for each of the groups (Motorized shoes: red; Split-belt: blue). The small bar plots on the right indicate the mean ± standard errors for the step lengths for each group during medium baseline.

### Smaller speed difference with the motorized shoes reduced the adaptation of StepPosition

We observed between-group differences in the adaptation of StepPosition (quantifying spatial asymmetry), but not StepTime (quantifying temporal asymmetry). This was indicated by the significant group by epoch interaction found in StepPosition (F_(3,48)_=3.47, p=0.023), but not in StepTime (F_(3,48)_=2.39, p=0.09) (Figure 4). Post-hoc analyses indicated that these differences in StepPosition were driven by distinct early and late adaptation values of this parameter in the Motorized shoes group compared to the Split-belt group (early adaptation: p = 0.031; late adaptation: p = 0.036). Yet, after-effects in StepPosition were significant in both groups (Motorized shoes: p < 0.001; Split-belt: p < 0.001) and after-effects in StepTime were only significant in the Motorized shoes group (Motorized shoes: p = 0.017; Split-belt: p =0.087) Interestingly, we also found a group effect (F_(1,48)_ = 6.58, p = 0.021) on StepVelocity and a group by epoch interaction trending effect (F_(1,48)_ = 2.78, p = 0.051) (Figure 4C). In particular, the StepVelocity was smaller in the group with Motorized shoes than in the Split-belt group during late adaptation (p=0.001), which we thought could impact the motor adaptation of the Motorized shoes group. Thus, we performed a correlation analysis on the late adaptation epoch with either StepTime or StepPosition as the dependent variable and StepVelocity as the predictor. We indeed found that larger speed differences between steps (i.e., larger StepVelocity values) were associated with higher steady-state values for both StepPosition (r= −0.52; p=0.032) and StepTime (r= −0.76; p<0.001) (Figure 4D). Moreover, the inter-subject variability in steady-state values was associated to individual after-effects in StepPosition (r=0.57; p=0.017), but not StepTime (r= −0.052; p=0.84) (Figure 4E). To sum up, the reduced speed difference in the Motorized shoes group limited the adaptation of StepPosition, but we still observed group after-effects with the motorized shoes in the spatial and temporal domains.

**Figure 4.**
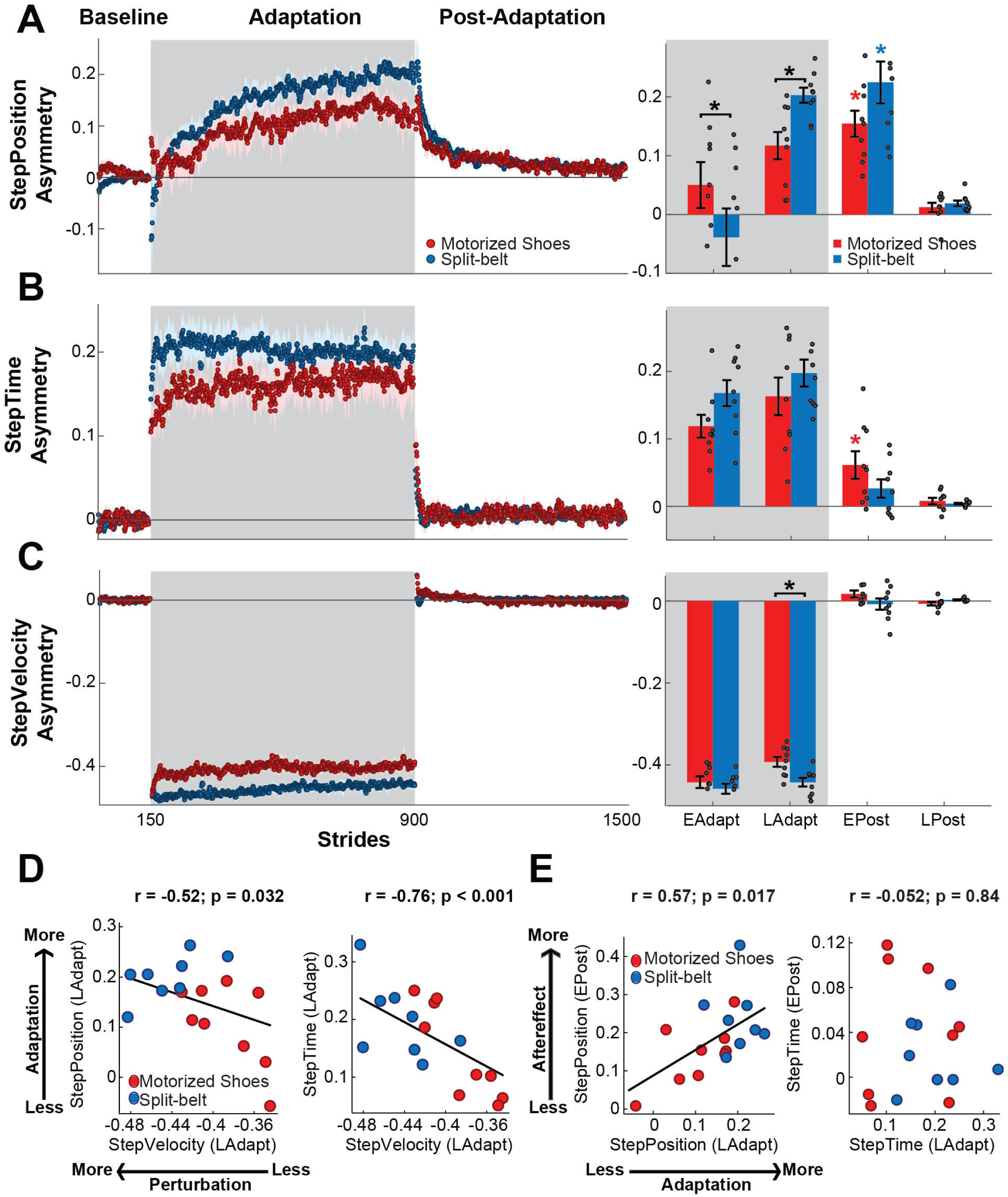
Adaptation of spatiotemporal components of step length asymmetry. (A, B, C- Left Panel) Time courses for StepPosition, StepTime, and StepVelocity before, during and after adaptation. Shaded gray areas represent the adaptation period during which the feet are moved at different speeds. “Tied” walking, both feet moved at the same speed, is indicated by the white regions. Colored dots represent the group average of 5 consecutive strides and colored shaded regions indicate the standard error for each group (Motorized shoes: red; Split-belt: blue). (A, B, C- Right Panel) The bar plots indicate the mean ± standard errors for StepPosition, StepTime, and StepVelocity for each group and epoch of interest. Gray dots represent individual subjects. Note that the values were corrected for baseline biases. Significant differences for post-hoc tests were indicated as follows. Black asterisks over the bracket above each epoch represent statistical significant differences between the Motorized shoes and the Split-belt groups (p<0.05). Colored asterisks over the bars indicate significant after-effects (i.e., early post-adaptation is significantly different from baseline; p<0.05) for each of the groups (Motorized shoes: red; Split-belt: blue). D) Scatter plots illustrate the association between the StepVelocity and the StepPosition and StepTime at steady-state during adaptation (i.e., LAdapt). Significant relations were observed between StepVelocity steady-state and StepPosition as well as StepTime steady-states. E) Scatter plots illustrate the association between the LAdapt and EPost for StepPosition and StepTime. Significant relations were observed for StepPosition, but not for StepTime.

### Similar cadence is observed between the groups throughout the experiment

We found that the motorized shoes did not alter the modulation of cadence throughout the experiment compare to split-belt walking (Figure 5 - left). Specifically, there were no significant group (F_(1,48)_=0.02, p=0.88) or group by epoch interaction effects on cadence (F_(3,48)_=0.32, p=0.81), indicating that the adaptation and after-effects of cadence were similar between groups (Figure 5 - right). We also found that both groups exhibited increased cadences during early post-adaptation compared to baseline (Motorized shoes: p = 0.002; Split-belt: p = 0.003). In sum, the motorized shoes modulate cadence similarly to the Split-belt group.

**Figure 5.**
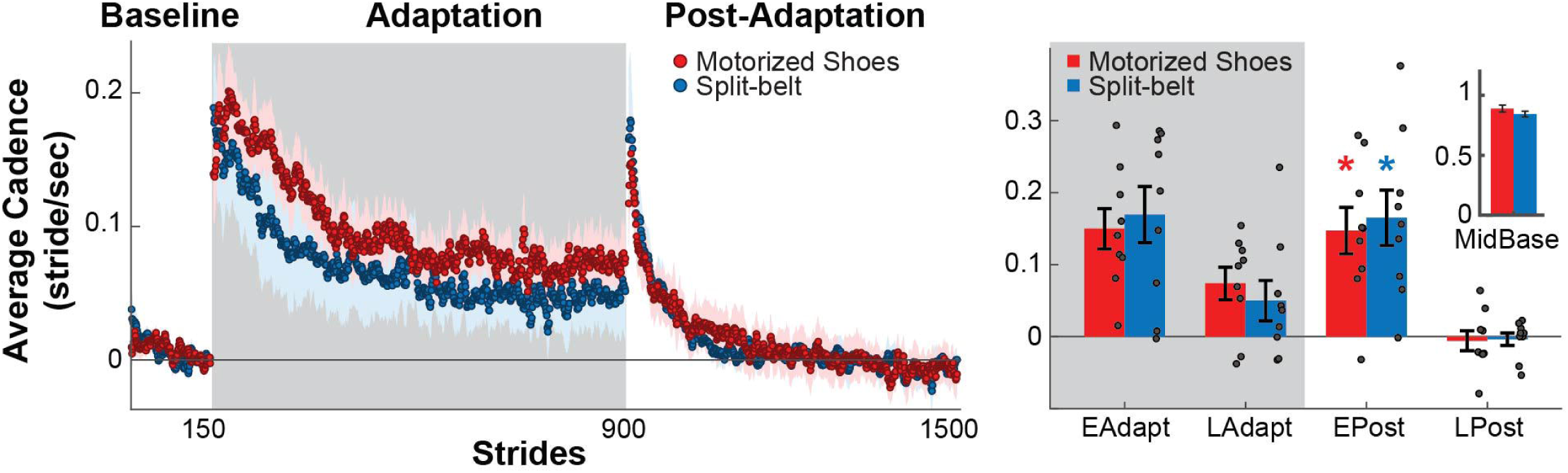
Modulation of cadence. (Left Panel) Time courses during medium baseline, adaptation and post-adaptation for the average cadence is shown for each group. Shaded gray areas represent the adaptation period during which the feet are moved at different speeds. “Tied” walking, both feet moved at the same speed, is indicated by the white regions. Colored dots represent the group average of 5 consecutive strides and colored shaded regions indicate the standard error for each group (Motorized shoes: red; Split-belt: blue). (Right Panel) Bar plots indicate the mean ± standard errors for cadence for each group and epoch of interest. Note that the values were corrected for baseline biases (i.e., MidBase). Colored asterisks over the bars indicate significant after-effects (i.e., early post-adaptation is significantly different from baseline; p<0.05) for each of the groups (Motorized shoes: red; Split-belt: blue). The small bar plot on the right indicate the mean ± standard errors for the Cadence for each group during medium baseline.

### Minimal effect of wearing motorized shoes on gait kinematics

Our result revealed a near-normal gait pattern in subjects walking with the motorized shoes. Figure 6A illustrates the joint angles over the gait cycle for the ankle, knee, and hip joints for the group wearing the motorized shoes (red) and the group wearing regular shoes (blue) during medium baseline walking (gray). We found joint angles were the same between groups for most phases of the gait cycle, in which significance was determined with an FDR controlling procedure (p>Pthreshold, Pthreshold = 0.0055, see methods) (Figure 6A). However, minor differences in joint angles were observed for specific gait cycle phases (18 comparisons, p<Pthreshold, Pthreshold = 0.0055). Specifically, the Motorized shoes group demonstrated reduced ankle dorsiflexion following ipsilateral heel strike and during late swing (double support DS1: p=0.004, effect size = 3.3°; late swing SW2: p=0.004, effect size = 4.1°). Moreover, the Motorized shoes group exhibited reduced knee flexion compared to the Split-belt group during early swing (SW1: p=0.004, effect size = 7.8°), followed by slightly more knee extension in late swing (SW2: p=0.001, effect size = 9.6°). Lastly, the Motorized shoes group had larger hip flexion during stance of baseline walking (p = 0.005, effect size = 4.1°). These group differences in baseline joint kinematics might be due to the additional weight of the motorized shoes (Ochsmann et al., 2016). In addition to baseline joint kinematics, we also compared late adaptation kinematics across groups (Figure 6B). Specifically, we contrasted the changes in joint angles during late adaptation relative to the speed-specific baseline for each of the six phases of the gait cycle. We found no differences between the groups (36 comparisons, p>Pthreshold), suggesting that joint angles were modulated similarly in the split condition with the motorized shoes or the split-belt treadmill. Thus, our results demonstrated that walking with the motorized shoes had only minor effects on joint kinematics and did not alter the adaptation of individual joint angles during split walking.

**Figure 6.**
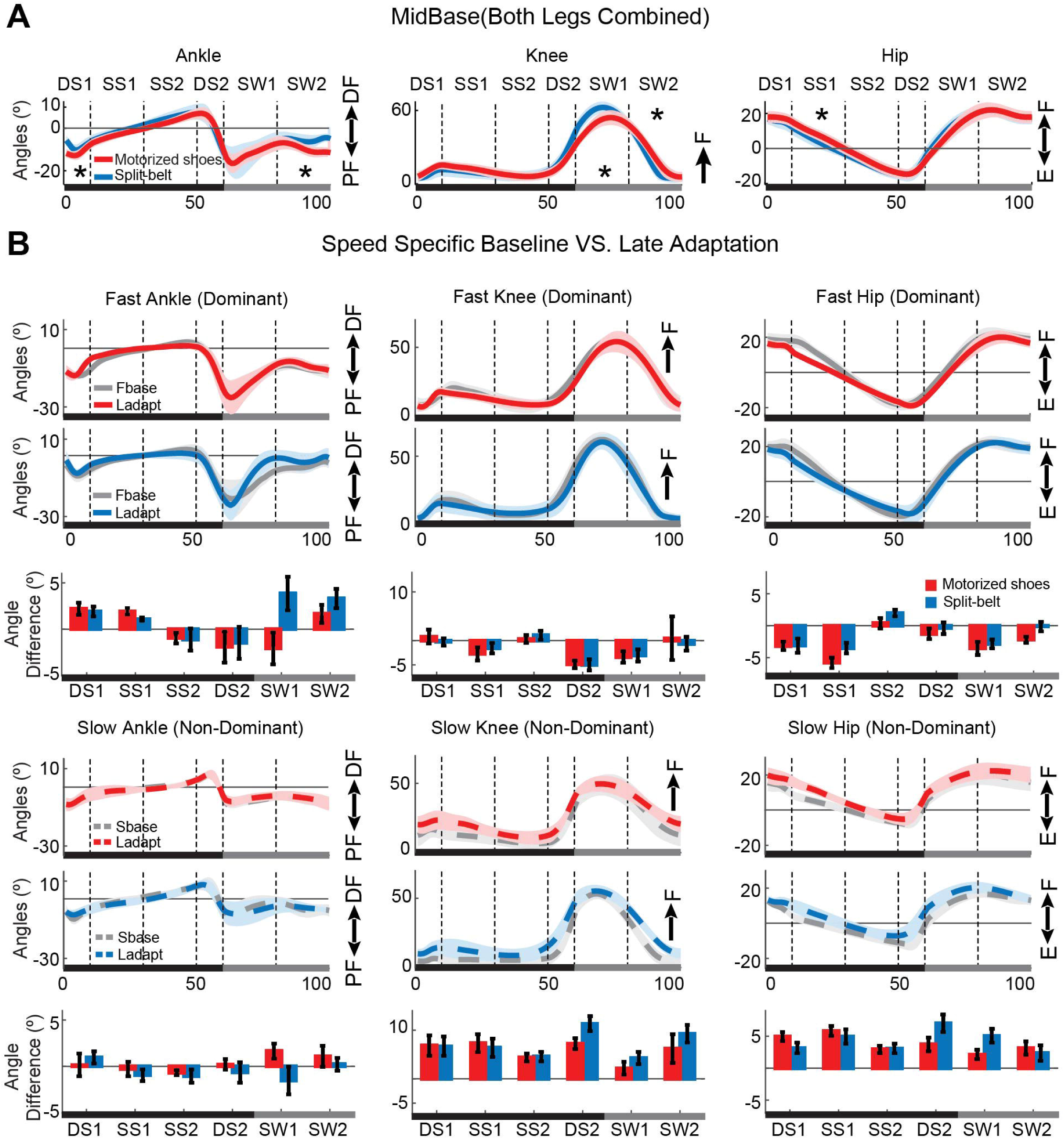
Joint angles during baseline and adaptation. *A)* Medium baseline joint angle trajectories with and without the motorized shoes. Solid lines represent the group average for the Motorized shoes (red) and the Split-belt groups (blue). Shaded areas represent standard errors. Asterisks indicate instances during the gait cycle when joint angles were significantly different across groups. The overall motion for all joints was similar across groups, but hip flexion, knee flexion and ankle dorsiflexion were smaller when wearing the motorized shoes. *B)* Speed specific baseline (gray) and steady-state angle trajectories during adaptation for the Motorized shoes (red) and the Split-belt (blue) groups. Solid lines represent the motion of the leg walking fast in the split condition (colored lines) and in the fast baseline (gray) condition. The dashed lines represent the motion of the leg walking slow in the split condition (colored lines) and in the slow baseline (gray) condition. The bars represent the change from the speed specific baseline to late adaptation in joint angles during different phases of the gait cycle. DS: Double support; SS: Single Stance; SW: Swing; DF: dorsiflexion; PF: plantarflexion; F: flexion; E: extension.

## 4 Discussion

### Summary

We investigated if a pair of motorized shoes could induce split-like locomotor adaptation. We found that the adaptation effects induced by the motorized shoes moving at different speeds were as robust as those observed with a split-belt treadmill. Moreover, we found that the gait pattern was largely similar between walking with the motorized shoes or on the split-belt treadmill. Specifically, the step length asymmetry, the cadence, and the step lengths were similar across groups during and after the split condition with either device. We only observed subtle differences in individual joint angles during the baseline condition with the motorized shoes compared to walking with regular shoes, which might be due to the greater height and weight of the motorized shoes. Taken together, our results suggest motorized shoes can induce robust sensorimotor adaptation in locomotion, opening the exciting possibility to study locomotor learning under more realistic situations outside the laboratory setting.

### Similar walking and adaptation with split-belt treadmill and with motorized shoes

We demonstrated that the motorized shoes can induce locomotor adaptation largely similar to the adaptation induced with the split-belt treadmill. This was shown by the comparable adaptation across groups of gait parameters, such as step length asymmetry, and the same modulation of joint angles from baseline to adaptation for both groups. Namely, the initial and steady state values during the split condition for the split-belt group and motorized shoes group were consistent with values previously reported for joint angle kinematics (Reisman et al., 2005; Winter, 1987) and asymmetries in step length (Finley et al., 2015; Malone and Bastian, 2010), step position (Sombric et al., 2017), and step time (Gonzalez-Rubio et al., 2019). In contrast, we found between-group differences in the fast step length during early adaptation, which were due to distinct placement of the leading foot when stepping with the motorized shoes compared to the regular shoes. In other words, subjects with the motorized shoes placed the foot closer to the body than with regular shoes. This distinct behavior might be explained by the fact that the balance is perturbed in the beginning of the split condition (Buurke et al., 2018; Iturralde and Torres-Oviedo, 2019) and it might be further challenged when stepping with the motorized shoes by augmenting the center of mass’ height, increasing even further gait instabilities while walking.

Of note, subjects with the motorized shoes reached lower steady state values of StepPosition (spatial) and slightly lower steady state values of StepTime (temporal) relative to the split-belt group. This was associated to the smaller speed differences that the Motorized shoes group experienced compared to the split-belt group, as indicated by our regression analysis. Thus, perturbation size regulated the extent to which subjects adapted, as observed in other sensorimotor adaptation protocols of reaching (Marinovic et al., 2017; Morehead et al., 2015) or walking (Finley et al., 2015; Yokoyama et al., 2018). Despite the subtle differences during adaptation, we saw similar after-effects between groups during early post-adaptation in all gait parameters. For example, cadence exhibited comparable changes between the groups during early adaptation and early de-adaptation, which is consistent with previous literature showing that stride time (i.e., inversely related to cadence) decreases in the beginning of adaptation (Reisman et al., 2005) and post-adaptation (MacLellan et al., 2014). In summary, our portable devised induced significant adaptation and after-effects of gait asymmetries in space and time opening the door for studying locomotor adaptation outside of the laboratory.

We also found a direct correspondence between adaptation and after-effects in the spatial domain, but not the temporal one. In particular, after-effects were positively associated to steady-states in StepPosition: the larger the steady-state value (relative to baseline), the more after-effects. This positive relation between steady state values and after-effects is commonly found in reaching or saccadic movements with well-defined performance errors (Chen-Harris et al., 2008) and spatial gait parameters in walking (Green et al., 2010; Sombric et al., 2019). However, this direct relation between steady state values during Adaptation period and the after-effects is more elusive when considering the temporal control of the limb. More specifically, temporal parameters, such as StepTime asymmetry, can change dramatically during the Adaptation period (i.e., split condition) without showing any significant after-effects (Gonzalez-Rubio et al., 2019; Long et al., 2015). Thus, it was unexpected to observe significant after-effects in StepTime asymmetry with the motorized shoes. Taken together these findings further support the idea of dissociable neural structures mediating the adaptation of spatial and temporal gait features (Boyd and Winstein, 2004; Darmohray et al., 2019).

### Study Implications

We found a few differences in joint motions when walking with our motorized shoes during regular walking, which will be useful for future designs of this portable device. Notably, we observed gait changes during baseline walking (i.e., both feet moving at the same speed) with the motorized shoes that were consistent with other studies showing that shoe weight (Ochsmann et al., 2016) and height (McDonald et al., 2019) alter walking movements. In addition, the rigidity of the motorized shoes’ soles (Chiou et al., 2012)is another factor that might contribute to the differences that we observed in joint angles during baseline walking. Thus, our gait analysis enabled us to identify key features that we will modify to create more a naturalistic walking conditions with the motorized shoes. This is important because contextual differences when wearing the motorized shoes could limit the extent of generalization of movements from walking with them to walking without this portable device. Locomotor adaptation with the motorized shoes overground could certainly reduce context specific difference that limit the generalization of treadmill movements, such as visual flow (Torres-Oviedo and Bastian, 2012), walking speed (Dingwell et al., 2001), and step initiation. However, it remains to be determine whether contextual cues due to the height, weight, and rigidity of the motorized shoes would also limit the generalization of locomotor learning with them.

Nevertheless, our results are promising because of the portability and low-cost of our device allowing us to use them outside the laboratory setting. This is exciting because we will be able to study gait under more realistic situations, such as when walking with variable gait speeds. It is well-accepted that motor variability can impact motor learning (Ulman et al., 2019; Wu et al., 2014), and walking on a treadmill is less variable compared to overground walking (Dingwell et al., 2001). Thus, having a device that can induce locomotor adaptation overground would help us gain more understanding about the relationship between variability and motor adaptation in a naturalistic behavior, such as walking. Moreover, learning a new task involves generation of new neural activity patterns, which appears after several days of practice (Oby et al., 2019). Our device will enable training over longer periods of time because individuals will be able to train at home and gain much more practice in the altered split environment than what is currently available. This can help us contribute to recent efforts to investigate the effect of long-term practice (Hardwick et al., 2019).

There have been efforts to develop portable rehabilitation devices (Afzal et al., 2015; Calabrò et al., 2018; Handzic et al., 2011; Lahiff et al., 2016) and assistive devices (Awad et al., 2017; Bae et al., 2018; Rao et al., 2008) to improve walking patterns in individuals with gait asymmetries, such as individuals post-stroke. While these apparatus could reduce the metabolic cost associated to gait in this clinical population (Awad et al., 2017) and improve walking speed (Buesing et al., 2015; Calabrò et al., 2018; Rao et al., 2008), these devices were unsuccessful in modifying the step length asymmetry (Handzic et al., 2011), which is an important parameter in rehabilitation of post-stroke patients (Patterson et al., 2008, 2014). For example, Lahiff and colleagues were able to modify push-off and breaking forces, but their device was unable to change step length of the participants (Lahiff et al., 2016). Similarly, Handzic and colleagues designed a device to passively induce a speed difference between the feet (Handzic et al., 2011; Handzic and Reed, 2013). However, this passive device induced limited changes in step length asymmetry post-adaptation (i.e., ∼ 5% of the after-effect size observed with the split-belt treadmill and motorized shoes). In sum, our study indicates that motorized shoes could tackle previous limitations altering gait asymmetries with portable devices and thus, could be potentially used to correct asymmetric steps post-stroke.

## 5 Author’s contribution statement

YA contributions include acquisition, analysis, and interpretation of the data, drafting the work and agreement to be accountable for all aspects of the work. XZ and RS contributions include development of the motorized shoes and providing technical expertise for using the motorized shoes. GT contributions include conception and design of the work, revising the work and agreement to be accountable for all aspects of the work. All authors contributed to revising the manuscript and providing a final approval of the version to be published.

